# The y-ome defines the thirty-four percent of *Escherichia coli* genes that lack experimental evidence of function

**DOI:** 10.1101/328591

**Authors:** Sankha Ghatak, Zachary A. King, Anand Sastry, Bernhard O. Palsson

## Abstract

Experimental studies of *Escherichia coli* K-12 MG1655 often implicate poorly annotated genes in cellular phenotypes. However, we lack a systematic understanding of these genes. How many are there? What information *is* available for them? And what features do they share that could explain the gap in our understanding? Efforts to build predictive, whole-cell models of *E. coli* inevitably face this knowledge gap. We approached these questions systematically by assembling annotations from the knowledge bases EcoCyc, EcoGene, UniProt, RefSeq, and RegulonDB. We identified the genes that lack direct experimental evidence of function (the “y-ome”) which include 1563 of 4653 unique genes (34%), of which 131 have absolutely no evidence of function. An additional 304 genes (6.6%) are pseudogenes or phantom genes. y-ome genes tend to have lower expression levels and are enriched in the termination region of the *E. coli* chromosome. Where evidence is available for y-ome genes, it most often points to them being membrane proteins and transporters. We resolve the misconception that a gene in *E. coli* whose primary name starts with “y” is unannotated, and we discuss the value of the y-ome for systematic improvement of *E. coli* knowledge bases and its extension to other organisms.

## Introduction

Unannotated genes in model organisms still play important roles in determining cell phenotype. This point was driven home by recent efforts to synthesize a minimal bacterial genome. The resulting syn3.0 organism includes just 473 genes, all of which are essential for growth, and a full 30% of which lack functional annotation (1,2). Even in *E. coli* K-12 MG1655, perhaps the best-studied model organism, unannotated genes often appear in experimental studies of strain engineering (3), laboratory evolution (4), and pathogenicity (5). These unannotated genes are one type of “dark matter” in the cell (6,7), and efforts to build predictive models of the genotype-phenotype relationship for whole cells will be hindered by unannotated genes that still affect cell phenotype (8,9).

Historically, unannotated genes in *E. coli* are known as “y-genes” because they have primary names starting with “y” (10)—not to be confused with “Y genes” which can indicate genes on the human Y chromosome (11). However, genes with primary names that begin with “y” are often functionally annotated. For example, in a recent study where *E. coli* was engineered to produce fatty acids via reversal of the fatty-acid beta-oxidation pathway, the authors knocked out the genes *yqeF* and *yqhD* to increase production of target molecules (3) and included the genes *ydiQRST, ydiO,* and *ydbK* in a predictive model of the cell (12). Searching for these genes in public knowledge bases such as EcoCyc (13) reveals that they vary greatly in annotation quality. Some (e.g., *yqhD)* are well-annotated with direct experimental evidence, while others (e.g., ydiO) have limited functional information. The variation of annotation quality between y-genes suggests a systematic approach to understanding the unannotated genes in *E. coli* is needed that goes beyond the primary gene name.

There are several model organism knowledge bases that represent the collected knowledge of the *E. coli* K-12 MG1655 genome: EcoCyc (13), EcoGene (14), UniProt (15), and RefSeq (16). Other useful knowledge bases cater to specific classes of gene products, such as the RegulonDB, which contains manually curated functional information about transcription factors in *E. coli* (17). Our initial review of these knowledge bases yielded conflicting information on gene function and level of annotation for many *E. coli* genes. Any attempt to systematically assess the function of unannotated genes must therefore draw from multiple knowledge bases and resolve these conflicts.

Many research groups have categorized *E. coli* genes and proteins by annotation quality as a part of their studies. In 2009, Hu *et al.* constructed a global functional atlas of *E. coli* proteins (18). First, they identified all unannotated proteins in the K-12 W3110 and MG1655 genomes. In order for a protein-encoding gene to be considered functionally uncharacterized in their analysis, it had to meet the following criteria: (i) The gene name begins with “y”, (ii) the gene does not have a known pathway within EcoCyc, and (iii) the gene does not have a functional description in GenProtEC (19) (any gene with a description containing the words “predicted”, “hypothetical”, or “conserved”). Based on these criteria, it was determined that 1431 of 4225 protein coding sequences were functionally unannotated. In 2015, Kim *et al.* published a database called EcoliNet that curated and predicted cofunctional gene networks for every protein coding gene in the *E. coli* genome (20). This study also quantified the number of uncharacterized protein coding genes in *E. coli.* To assess functional annotation, they used the presence of experimentally supported “biological process” annotations in the Gene Ontology database (21). They concluded that ~2000 protein coding genes in *E. coli* were functionally unannotated. The most comprehensive effort to assess the level of annotation in bacterial genomes has been Computational Bridges to Experiments (COMBREX) (22,23). The COMBREX knowledge base currently contains information about 4182 protein coding genes in *E. coli* K-12 MG1655, of which 2378 (57%) have experimentally verified function, 1741 (42%) have predicted but not experimentally verified function, and 63 (2%) have no predicted function. These studies of unannotated genes in *E. coli* K-12 MG1655 provided inspiration for this work. However, our effort covers both protein-coding and non-protein-coding genes, disregards nomenclature (i.e., whether a gene name begins with “y”) as an indicator of annotation quality, and is presented as a reproducible workflow to keep the analysis up-to-date as knowledge bases improve.

It is difficult to define a Y-ome because there are no established rules that specify the level of annotation necessary for a gene to be “well-annotated”. We draw on experience in systems biology to precisely define the Y-ome. The contribution of any gene function to cellular phenotype can now be codified computationally for various cellular systems, including metabolism (24), cell signaling (25), gene expression (26), and replication (9). Therefore, we define functional annotation as the information necessary to have an effect on phenotype predictions in a systems biology model. This allows us to define annotation differently for different cellular systems; for example, a metabolic enzyme should be annotated through enzymatic activity assay, a transcriptional regulator through DNA-binding assay and study of gene regulation. With this approach, we can be sure that the definition of annotation follows from the actual impact that the gene could have on cell phenotype. Such definitions are expected to evolve as cellular modeling methods evolve. Thus, the concept of a Y-ome can keep pace with new developments in the field.

To determine the Y-ome for *E. coli* K-12 MG1655, we first define it in precise terms. Next, we present a reproducible workflow to compile annotations from *E. coli* knowledge bases to determine the Y-ome. This workflow included an automated portion and a manual curation step to resolve 292 genes that could not be automatically assigned to a category. The resulting Y-ome includes 34% of *E. coli* genes. We describe some trends for these Y-ome genes, including their enrichment in the termination region of the *E. coli* chromosome, lower average expression levels than well-annotated genes, and evidence that certain types of genes (e.g., transporters) are more highly represented. Finally, we resolve the misconception that a gene in *E. coli* whose primary name starts with “y” is necessarily unannotated.

## Results & Discussion

### Definition of the Y-ome

The Y-ome encompasses all genes in an organism that lack functional annotation. However, it is difficult to precisely define whether a functional annotation is sufficient to call the gene “well-annotated.” We drew on systems biology to precisely define this boundary. In systems biology, predictive models can be used to link genotype to phenotype. In these models, the definition of functional annotation is that a gene can be mechanistically linked through a network to a measurable phenotypic effect. Thus, we define the Y-ome as *the set of genes that lack a known mechanistic effect on cell phenotype.* For a given gene, we can use the following test, “Is this gene sufficiently annotated that it can (1) be incorporated into a predictive model and (2) have an impact on the predicted phenotype?” The model in this test can be hypothetical, but it helps frame the precise definition of the Y-ome. Because genotype-phenotype models can take different forms (metabolic models have a different theoretical basis from regulatory models), this definition allows for different kinds of evidence based on the gene function (enzymatic assays for metabolic genes; DNA-binding assays and gene knockout studies for regulatory genes).

Our definition of the Y-ome also restricts annotations to experiments on the organism of interest and excludes annotations that are drawn entirely from gene sequence similarity. Sequence similarity provides evidence of gene function, but generalist proteins can play different roles depending on the cell context (27,28). Therefore, we looked for direct experimental evidence of function with the target organism or multiple lines of evidence beyond just sequence similarity. We also made an exception for insertion elements—these were considered well-annotated based on sequence similarity alone.

Genes that are well-annotated (not in the Y-ome) could potentially have secondary functions in the cell (29,30), so these genes could still require additional study to improve our understanding of the system, and model-driven approaches can be used to systematically identify them (28). Finally, genes with high sequence similarity might be pseudogenes or cryptic genes. Because they are not expected to affect cell phenotype, we excluded pseudogenes and cryptic genes from the Y-ome in this study. However, if they are found to contribute to cellular phenotype, they should be included.

### A workflow for identifying the *E. coli* Y-ome

To systematically determine the Y-ome for *E. coli* K-12 MG1655, we developed a semi-automated approach (Fig. 1) to identify unique genes across five *E. coli* knowledge bases and integrate their data to define a consensus Y-ome. The automated part of this process proceeded in three steps: (1) downloading data from each knowledge base, (2) extracting text-based features (Supplementary Data S2), and (3) using keywords to automatically assign each gene annotation in a knowledge base to the categories “Y-ome,” “Well-Annotated,” or “Not enough information for automated assignment” (Fig. 2a). Pseudogenes and cryptic genes were kept separate and marked “Excluded.” The keywords used to make these assignments are described in the Methods.

**Figure 1:**
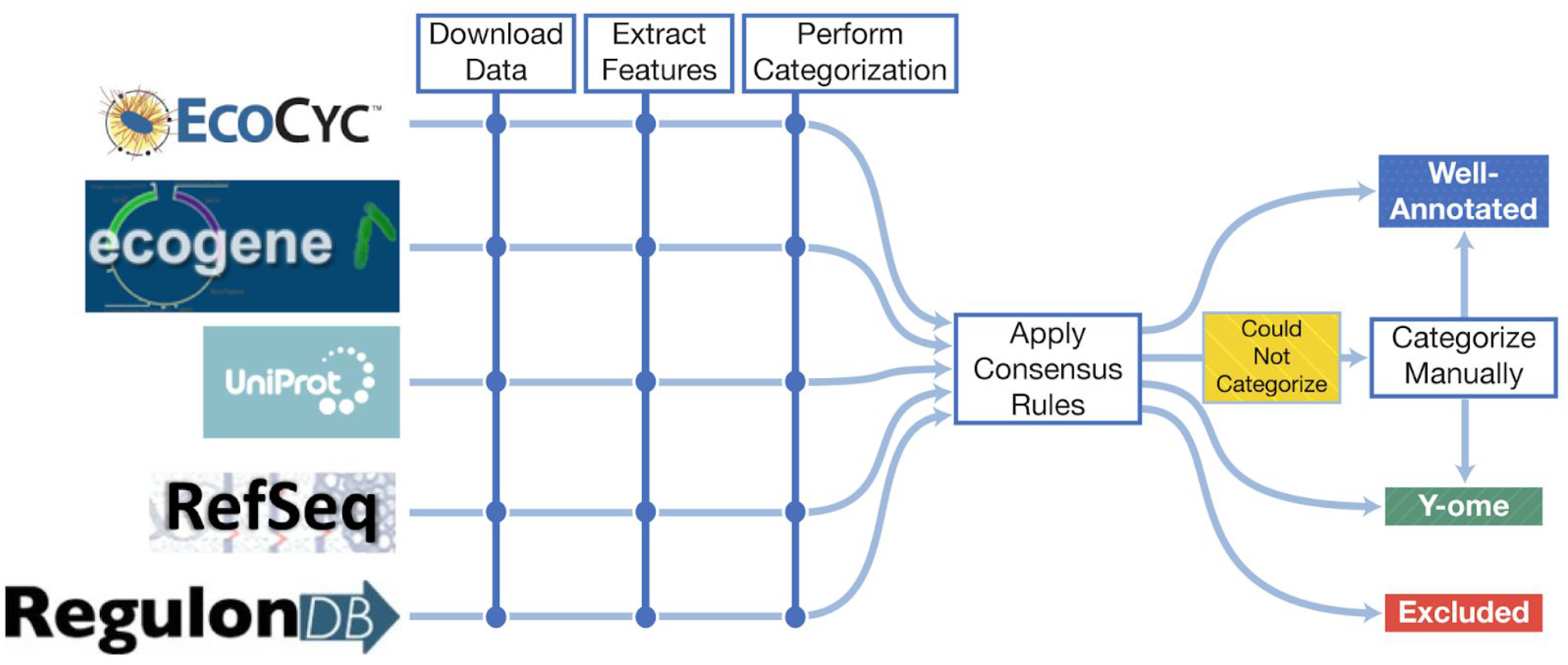
A workflow for defining the Y-ome of E. coli K-12 MG1655. Data was collected from five *E. coli* knowledge bases, and automated categorization was applied to determine their annotation level. Next, we applied a set of consensus rules to combine categorizations from multiple databases. When the consensus rules could not be applied, genes were manually curated and placed in one of the categories. Thus, genes were categorized as “Well-Annotated” or “Y-ome” according to the definition of the Y-ome (see Section “Definition of the Y-ome”). Pseudogenes and phantom genes were treated separately in the “Excluded” category.

**Figure 2:**
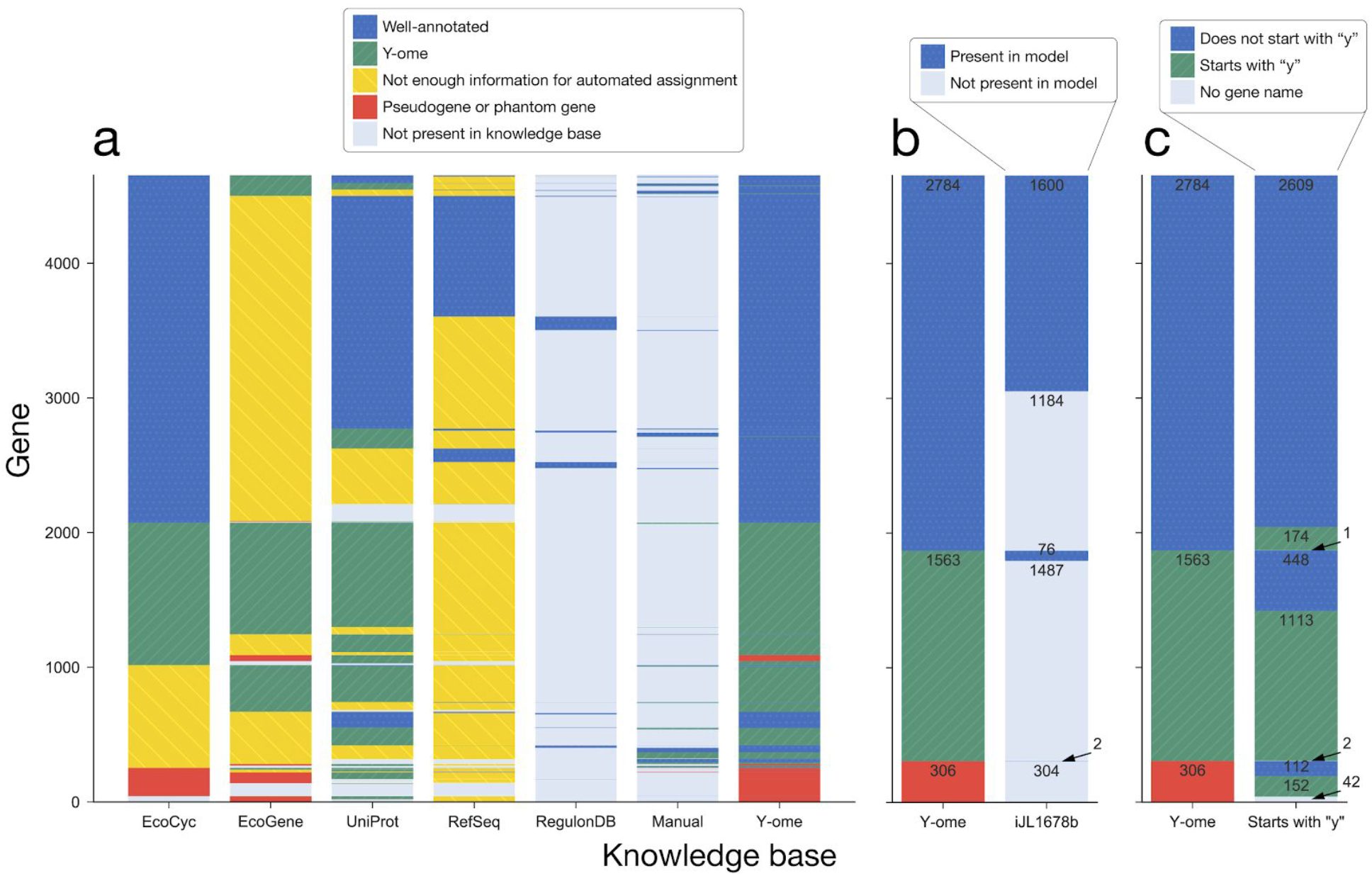
Gene annotation across knowledge bases. (a) An automated approach was used to categorize genes from each database as “Well-Annotated” or “Y-ome” based on the definition of the Y-ome. Pseudogenes and phantom genes were excluded. The resulting Y-ome includes 1563 genes. (b) Y-ome categories were compared to the content of the latest *E. coli* genome-scale ME-model. 1184 genes that are annotated and not in the ME-model represent an opportunity to increase the scope of whole-cell models. The 76 genes in the ME-model and in the Y-ome might have model-driven evidence of function that could be used to systematically annotate them. (c) 174 genes have primary names that start with “y” but are well-annotated, and 448 genes in the Y-ome have non-“y” primary names.

After assigning genes in each knowledge base to categories, consensus rules were applied to combine the results from the separate knowledge bases. In general, we checked first for an agreement among knowledge bases. For instance, if two knowledge bases indicated a gene was “Well-Annotated” and the others did not have enough information to assign a category, then the consensus was “Well-Annotated”. When databases disagreed, then no consensus was possible. In these cases, manual annotations were made (for 292 genes) based on reading the knowledge base entries and consulting the literature. Finally, certain kinds of structured evidence (e.g., the Experimental Evidence section in EcoCyc) were taken as high-quality evidence, and we ignored conflicting details from other knowledge bases in these cases. Pseudogenes and phantom genes were treated separately in the “Excluded” category, and for these genes we also ignored conflict between the knowledge bases. More detail on the consensus rules can also be found in the Methods.

Based on this analysis, we identified 4653 unique genes across *E. coli* K-12 MG1655 knowledge bases, and each was assigned to the “Y-ome”, “Well-Annotated”, or “Excluded” categories (Supplementary Data S1). Of these 4653 genes, 2784 have information that indicate a sufficient level of functional and regulatory evidence to exclude them from the Y-ome (Fig. 2a). Thus, 1563 genes (34%) are in the Y-ome of *E. coli* K-12 MG1655. No individual knowledge base provides information to fully define the Y-ome, but EcoCyc comes the closest. Of the 1563 Y-ome genes, there were 131 for which we found no information in the knowledge bases (see Methods) and 306 that were marked as pseudogenes or cryptic genes.

### Gene expression and chromosome Location

It was previously observed by Hu *et al.* that poorly annotated genes tend to be expressed at a lower level than well-annotated genes (18). We confirmed this with the Y-ome by comparing gene expression of Y-ome genes and well-annotated genes in a compendium of RNA-seq data from our research group. The dataset includes expression values for 4319 *E. coli* genes across 85 conditions. Genes in the Y-ome tend to have lower expression across the surveyed conditions (Fig. 3) with average normalized expression count for Y-ome genes being 234 compared to 1583 FPKM for well-annotated genes (t-test p-value < 1×10^-6^). Attempts to annotate Y-ome genes may be more successful if they prioritize the highly expressed Y-ome genes that have a greater potential to affect observable phenotypes. Alternatively, experiments that identify growth conditions with greater expression of Y-ome genes could help elucidate their functions because the genes are more likely to have a phenotypic effect under conditions where they are expressed.

**Figure 3:**
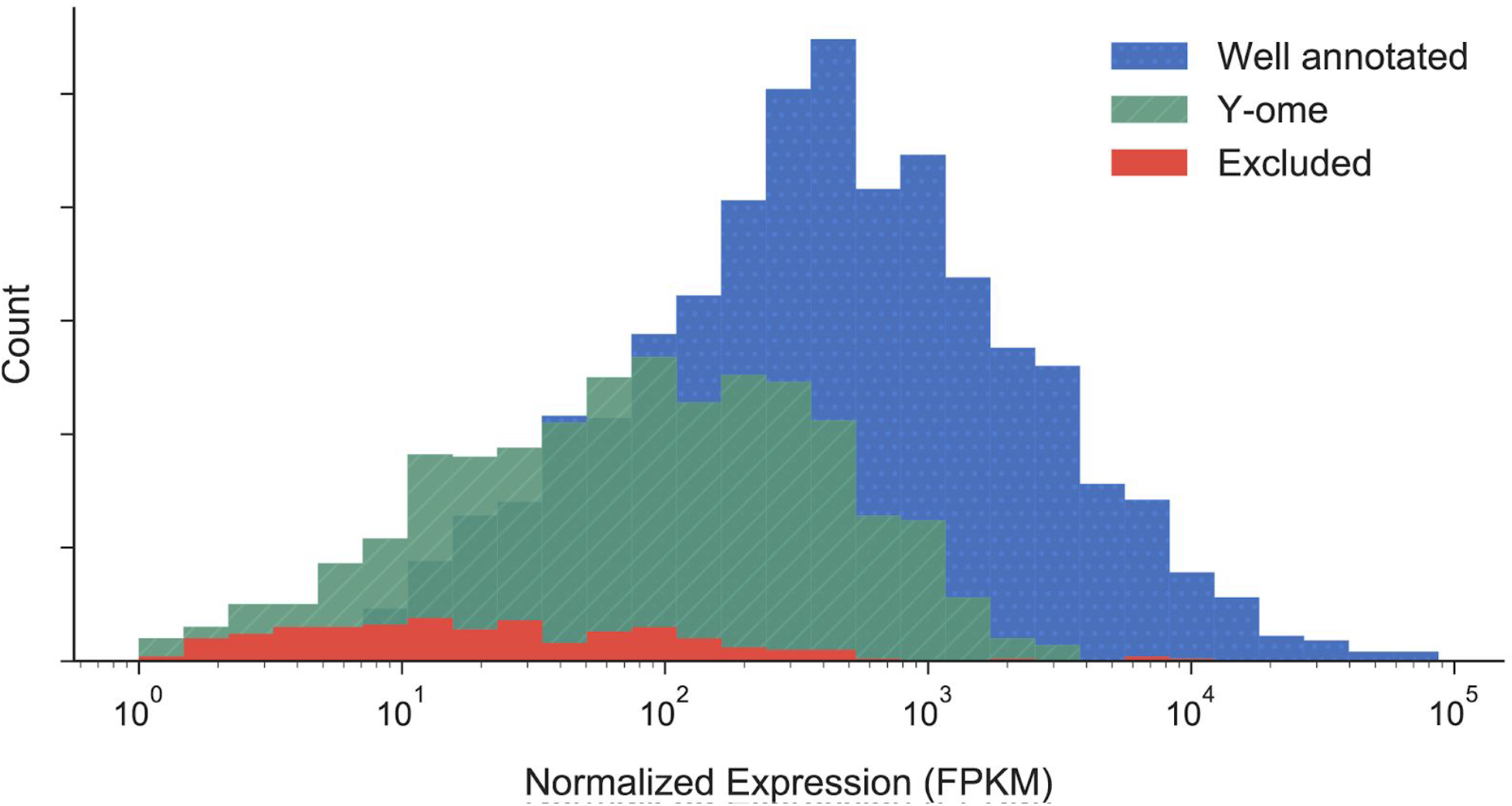
Average gene expression for all genes in a compendium of E. coli RNA-seq data. Histogram of normalized mean expression levels (FPKM) for Y-ome (green) and well-annotated (blue) genes across the 86 conditions surveyed in a compendium of RNA-Seq data.

We observed a low density of Y-ome genes near the origin of replication (ORI) of the *E. coli* chromosome and a high density of Y-ome genes in the termination region (opposite ORI, Fig. 4a). Highly expressed genes are known to be enriched near ORI (31–33), which was observed in our gene expression compendium (Fig. 4b). These observations tell a simple story of highly expressed genes that have obvious effects on phenotypes under laboratory conditions and are therefore well-annotated, and lowly expressed genes that do not affect phenotypes enough to be easily characterized. However, the Y-ome genes with highest expression (above an arbitrary cutoff of 1000 FPKM) are split between the origin and termination regions (Fig. 4a), which suggests that some other factor might be keeping genes near the termination region from being characterized. High-throughput gene annotation might shed further light on this phenomenon.

**Figure 4:**
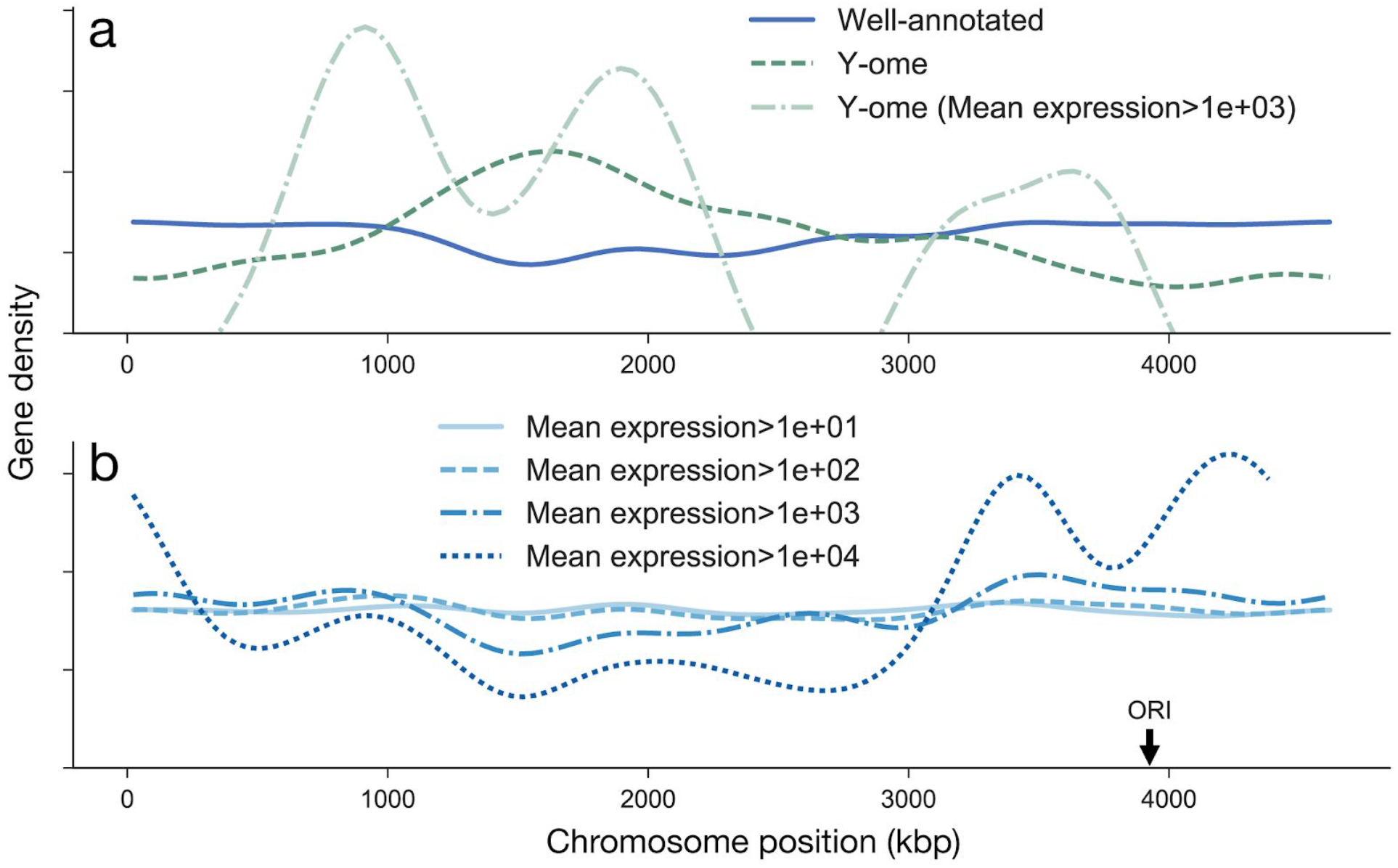
Gene expression by location on the chromosome. (a) Y-ome genes are enriched in the termination region of the *E. coli* chromosome, opposite the origin of replication (ORI). (b) Highly expressed genes are known to be enriched around the origin of replication (18), which we confirmed by plotting density of genes in the chromosome with increasing thresholds of mean gene expression (FPKM) across a compendium of RNA-seq data for 86 conditions.

### Functions of Y-ome genes

The Y-ome of *E. coli* provides clues to the function of Y-ome genes. The most common terms associated with Y-ome genes can easily be extracted from *E. coli* knowledge bases (Table 1). These terms indicate that many membrane-associated proteins (485 genes) and particularly transporters (284 genes) remain to be annotated. Membrane-bound proteins and transporters are particularly hard to characterize with certainty (34), but high-throughput methods might change that, as they already have for enzymatic assays (35), gene-environment networks (36), and protein-protein interactions (18). Thus, the Y-ome offers a set of candidate transport-associated genes for high-throughput analysis. High-throughput analysis could also be relevant for gene sets related to enzymes (269 genes), signaling (261 genes), lipoproteins (100 genes), and biofilms (69 genes). As evidence accumulates in *E. coli* knowledge bases, this workflow can be rerun to improve the candidate gene sets.

**Table 1:** The most common words found in knowledge bases features for Y-ome genes. The counts indicate the number of unique Y-ome genes for which each phrase appears. Similar words are grouped into sets.

| Word | Y-ome<br>(n=1563) | Well-Annotated<br>(n=2784) | Excluded<br>(n=306) |
| --- | --- | --- | --- |
| peptide/polypeptide/protein(s) | 1533 | 2535 | 217 |
| inner/outer/membrane/transmembrane | 485 | 744 | 41 |
| binds/binding | 421 | 1475 | 25 |
| regulate(s)/regulated/regulator(y)/regulation/regulon | 367 | 884 | 33 |
| transport/transporter/export/import | 284 | 676 | 53 |
| enzyme | 269 | 1576 | 15 |
| signal | 261 | 255 | 29 |
| initiation | 245 | 408 | 30 |
| phage/prophage | 185 | 181 | 69 |
| oxidoreductase/reductase/reduce | 141 | 356 | 6 |
| transcription/transcriptional | 129 | 374 | 12 |
| promoter | 118 | 198 | 4 |
| periplasm/periplasmic | 113 | 324 | 6 |
| lipoprotein | 100 | 86 | 5 |
| transferase | 93 | 374 | 19 |
| resistance | 91 | 240 | 1 |
| phosphate | 72 | 811 | 9 |
| synthesis | 70 | 730 | 4 |
| biofilm | 69 | 130 | 9 |
| lysis | 64 | 356 | 7 |
| metabolism | 63 | 412 | 8 |
| structural | 63 | 151 | 7 |
| sugar | 59 | 95 | 10 |
| stress | 58 | 182 | 1 |
| hydrogen | 56 | 260 | 3 |
| aerobic | 56 | 214 | 0 |

## Conclusion

The Y-ome represents a systematic accounting of genes lacking direct experimental evidence of function and should be valuable for guiding systematic gene function discovery. The accumulation of omics data is still accelerating, so a Big Data approach where multi-omic data types are combined with manual analysis and machine learning to derive new predictions shows promise towards elucidating the Y-ome (35,37). A data-driven discovery workflow could also be extended to poorly characterized organisms where high-throughput approaches to genome annotation can accelerate our accumulation of biological knowledge. The Y-ome is an ideal input to these approaches, and it can provide insights into possible gene function when paired with a genome annotation or gene expression data.

We defined the Y-ome by considering the effect a gene might have on a predictive model of cell phenotype, and this approach has corollary benefits for the longstanding effort to build predictive whole-cell models. Computational models are now being extended to include all cellular functions (8,38). The Y-ome encompasses genes that cannot be included in models because their contributions to cellular phenotypes are not understood (9). While whole-cell models provide a systematic approach to organizing our knowledge of organisms, the Y-ome is a systematic approach to evaluating our *lack* of knowledge in a genome. Comparing the 2784 well-annotated genes in *E. coli* to the 1678 genes in the latest genome-scale ME-model (39), it is clear that the models can grow by over a thousand genes before running up against our lack of knowledge (Fig. 2B). However, because unannotated genes are known to affect cell phenotype, the content of the Y-ome will have to eventually be addressed.

In 1998, a year after the first *E. coli* genome was released, Kenneth Rudd proposed a systematic naming scheme for unannotated open reading frames where each was given a unique name starting with the letter “y” (10). This is a convenient system, but the community did not settle on an official mechanism for assigning new names for these y-genes when functions were established. The tradition has been for newly identified functions to be published along with a proposed primary name, leaving it to peer reviewers to call out duplicate names and other issues. But without a central mechanism for standardized naming, many y-genes have been annotated without receiving new names (174 genes, Fig. 2c). And poorly annotated genes have received new names not starting with “y” because their function was partially established, determined based on computational predictions, or based on presence in an operon (448 genes, Fig. 2c). With the Y-ome, we can decouple gene names from assessments of functional annotation and provide a more consistent resource for anyone interested in systematic analysis of unannotated genes.

The concept of a Y-ome can be applied to any genome, and we hope that the provided workflow will inspire development of the Y-ome of other organisms. Knowledge bases that use the same knowledge base structure across organisms (e.g., EcoCyc/BioCyc) are a good place to start because many features of the workflow can be directly applied to a new organism. The developers and curators of knowledge bases play a central role in enabling this kind of workflow. To wit, the most useful feature across the five knowledge bases we analyzed was the “Evidence” section for EcoCyc genes; this section provides structured data on the experimental evidence for gene function, along with a literature reference. This feature is available now for the human genome on HumanCyc, so quick progress could made on the human Y-ome.

## Methods

### A workflow to determine the *E. coli* Y-ome

We collected data from the following five knowledge bases: EcoCyc release 21.5 (13), EcoGene version 3.0 (14), UniProt release 2017_11 (15), RefSeq *via* genome accession NC_000913.3 (16), and RegulonDB version 9.4 (17). We extracted features from the downloaded data, and we populated a relational (SQLite) database with them.

Definitions of pseudogenes and phantom genes were taken from both EcoGene and EcoCyc. In EcoGene, pseudogenes are indicated by a primary name ending in an apostrophe. EcoCyc explicitly defines lists of pseudogenes and phantom genes, currently available here: https://ecocyc.org/ECOLI/class-instances?object=Pseudo-Genes https://ecocyc.org/ECOLI/class-instances?object=Phantom-Genes

Keywords were used to automatically categorize genes for each knowledge base feature. To identify the keywords, we read the knowledge bases to look for commonly used phrases in the parlance of the particular knowledge base signified the level of annotation. For example, in EcoCyc, the keywords “possibly”, “predicted”, “hypothetical”, “putative”, “conserved”, “uncharacterized protein”, and “No information about this” were used to identify low annotation (Y-ome), and the keywords “assay”, “traceable author statement to experimental support”, and “reaction blocked in mutant” were used to identify genes with high annotation (Well-Annotated). The full list of keywords are available in our workflow in the “sources” directory.

Defining keywords is a subjective process, so we relied on structured data whenever possible. To determine annotation level for UniProt genes, we used the “annotation score” for each associated protein. Annotation scores of 2 or below were used to indicate “Y-ome” and 4 or above to indicate “Well-Annotated”. Genes with annotation score 3 were categorized as “Not enough information for automated assignment”.

### Consensus rules

After automatic categorization for individual knowledge bases, consensus rules were applied to generate a final Y-ome list. First, if categorizations from the knowledge bases were in agreement (e.g. all Y-ome or all Well-Annotated), then that annotation was chosen as the consensus. If no automatic annotation could be made (i.e. all knowledge bases had the category “Not enough information for automated assignment” for a gene) or if a conflict between annotations was identified, then the gene was marked for manual annotation. 292 genes were manually annotated—using the knowledge bases and the primary literature—to finalize the Y-ome list. There were four exceptions that were identified as heuristics to improve the quality of the final list:

1. The Evidence section for EcoCyc genes is a high-quality, manually-curated, and structured read-out of annotation quality based on the literature. Therefore, we gave this section priority over other data in the workflow. Particularly, we looked for Evidence features with keywords “assay”, “reaction blocked in mutant”, and “traceable author statement to experimental support” which were marked as Well-Annotated in the final categorization.
2. RegulonDB contains curated and experimentally-validated annotations of transcription factors. function. Thus, genes with “Strong” evidence in RegulonDB were marked as Well-Annotated in the final categorization.
3. When EcoCyc and UniProt were both categorized as Well-Annotated for a given gene, then this gene was automatically marked as Well-Annotated in the final categorization. This heuristic was helpful to identify cases where EcoGene was missing key evidence that the other knowledge bases had picked up (e.g. *dhaM).*
4. Insertion elements, identified in EcoCyc by gene names beginning with “ins”, were considered to be Well-Annotated in the final categorization.

### Genes with no information

To identify genes for which no information at all is available, we filtered the database for genes with features drawn from knowledge-base-specific phrase lists that corresponded to genes with no other functional information. For example, “Putative uncharacterized” often appeared in such UniProt entries. As another example, EcoCyc genes with no information have summaries that begin with this phrase “No information about this”. (e.g. “No information about this protein was found by a literature search conducted on February 23, 2017” for *ybiU).* The full list of phrases that were used can be found in the file “notebooks/17.11.02 Genes with no information.ipynb” in the workflow repository for this project (see Section Reproducibility).

When genes were annotated only with a protein domain or family, we still included them in the list because such domains (e.g. DUF1479 for ybiU) often themselves have no functional information associated (DUF1479 has the description “Protein of unknown function” on Pfam: https://pfam.xfam.org/family/PF07350).

### Gene expression compendium

A compendium of RNA-seq data for *E. coli* K-12 MG1655 (wild type, single gene mutants, and laboratory evolution endpoints) was used to analyze expression of Y-ome genes. All RNA-seq experiments were conducted using the protocol described by Seo *et al.* (40). A total of 85 unique strain-condition pairs are included in the compendium, and it will soon be available for public use. Fragments per kilobase of exon per million fragments (FPKM) were calculated using Cufflinks (41), and the mean FPKM across all 85 conditions was calculated to generate a comparison of gene expression between Y-ome genes, well-annotated genes, and excluded genes (Fig. 3).

### Chromosome location & density

Gene locations were extracted from gene start sites in the *E. coli* RefSeq genome annotation NC_000913.3. Gene density plots were created with a circular kernel density estimation method using the von Mises distribution.

### Reproducibility

All code and data necessary to reproduce this analysis can be found on GitHub and with a permanent DOI on Zenodo: https://github.com/zakandrewking/y-ome https://doi.org/10.5281/zenodo.1240561

## Acknowledgments

Special thanks to Marc Abrams for his help with editing, clarification, and discussion, to Ye Gao and Donghyuk Kim for providing feedback on the approach, and to Devleena Kole for her support and suggestions.

## References

1. Hutchison, C. A., 3rd, Chuang, R.-Y., Noskov, V. N., et al. (2016) Design and synthesis of a minimal bacterial genome, Science, 351, aad6253.

2. Danchin, A. and Fang, G. (2016) Unknown unknowns: essential genes in quest for function, Microb. Biotechnol., 9, 530–540.

3. Dellomonaco, C., Clomburg, J. M., Miller, E. N., et al. (2011) Engineered reversal of the *β*-oxidation cycle for the synthesis of fuels and chemicals, Nature, 476, 355–359.

4. Sandberg, T. E., Pedersen, M., LaCroix, R. A., et al. (2014) Evolution of Escherichia coli to 42 °C and subsequent genetic engineering reveals adaptive mechanisms and novel mutations, Mol. Biol. Evol., 31, 2647–2662.

5. Hufnagel, D. A., DePas, W. H. and Chapman, M. R. (2014) The disulfide bonding system suppresses CsgD-independent cellulose production in Escherichia coli, J. Bacteriol., 196, 3690–3699.

6. Maumus, F. and Quesneville, H. (2014) Deep investigation of Arabidopsis thaliana junk DNA reveals a continuum between repetitive elements and genomic dark matter, PLoS One, 9, e94101.

7. Johnson, J. M., Edwards, S., Shoemaker, D., et al. (2005) Dark matter in the genome: evidence of widespread transcription detected by microarray tiling experiments, Trends Genet., 21, 93–102.

8. Bordbar, A., Monk, J. M., King, Z. A., et al. (2014) Constraint-based models predict metabolic and associated cellular functions, Nat. Rev. Genet., 15, 107–120.

9. Karr, J. R., Sanghvi, J. C., Macklin, D. N., et al. (2012) A whole-cell computational model predicts phenotype from genotype, Cell, 150, 389–401.

10. Rudd, K. E. (1998) Linkage map of Escherichia coli K-12, edition 10: the physical map, Microbiol. Mol. Biol. Rev., 62, 985–1019.

11. Ballouz, S., Dobin, A., Gingeras, T. R., et al. (2018) The fractured landscape of RNA-seq alignment: the default in our STARs, Nucleic Acids Res.

12. Cintolesi, A., Clomburg, J. M. and Gonzalez, R. (2014) In silico assessment of the metabolic capabilities of an engineered functional reversal of the *β*-oxidation cycle for the synthesis of longer-chain (C≥4) products, Metab. Eng., 23, 100–115.

13. Keseler, I. M., Mackie, A., Peralta-Gil, M., et al. (2013) EcoCyc: Fusing model organism databases with systems biology, Nucleic Acids Res., 41, 605–612.

14. Zhou, J. and Rudd, K. E. (2013) EcoGene 3.0, Nucleic Acids Res., 41, D613–24.

15. UniProt Consortium (2015) UniProt: a hub for protein information, Nucleic Acids Res., 43, D204–12.

16. O’Leary, N. A., Wright, M. W., Brister, J. R., et al. (2016) Reference sequence (RefSeq) database at NCBI: current status, taxonomic expansion, and functional annotation, Nucleic Acids Res., 44, D733–45.

17. Gama-Castro, S., Salgado, H., Santos-Zavaleta, A., et al. (2016) RegulonDB version 9.0: high-level integration of gene regulation, coexpression, motif clustering and beyond, Nucleic Acids Res., 44, D133–43.

18. Hu, P., Janga, S. C., Babu, M., et al. (2009) Global functional atlas of Escherichia coli encompassing previously uncharacterized proteins, PLoS Biol., 7, e96.

19. Serres, M. H., Goswami, S. and Riley, M. (2004) GenProtEC: an updated and improved analysis of functions of Escherichia coli K-12 proteins, Nucleic Acids Res.

20. Kim, H., Shim, J. E., Shin, J., et al. (2015) EcoliNet: a database of cofunctional gene network for Escherichia coli, Database, 2015.

21. Gene Ontology Consortium (2015) Gene Ontology Consortium: going forward, Nucleic Acids Res., 43, D1049–56.

22. Anton, B. P., Chang, Y.-C., Brown, P., et al. (2013) The COMBREX project: design, methodology, and initial results, PLoS Biol., 11, e1001638.

23. Chang, Y.-C., Hu, Z., Rachlin, J., et al. (2015) COMBREX-DB: an experiment centered database of protein function: knowledge, predictions and knowledge gaps, Nucleic Acids Res., gkv1324.

24. Reed, J. L., Famili, I., Thiele, I., et al. (2006) Towards multidimensional genome annotation, Nat. Rev. Genet., 7, 130–141.

25. Papin, J. A. and Palsson, B. O. (2004) The JAK-STAT signaling network in the human B-cell: an extreme signaling pathway analysis, Biophys. J., 87, 37–46.

26. O’Brien, E. J., Lerman, J. A., Chang, R. L., et al. (2013) Genome-scale models of metabolism and gene expression extend and refine growth phenotype prediction, Mol. Syst. Biol., 9, 693.

27. Nam, H., Lewis, N. E., Lerman, J. a., et al. (2012) Network context and selection in the evolution to enzyme specificity, Science, 337, 1101–1104.

28. Guzmán, G. I., Utrilla, J., Nurk, S., et al. (2015) Model-driven discovery of underground metabolic functions in Escherichia coli, Proceedings of the National Academy of Sciences, 112, 929–934.

29. D’Ari, R. and Casadesús, J. (1998) Underground metabolism, Bioessays, 20, 181–186.

30. Notebaart, R. A., Szappanos, B., Kintses, B., et al. (2014) Network-level architecture and the evolutionary potential of underground metabolism, Proceedings of the National Academy of Sciences, 111, 11762–11767.

31. Allen, T. E., Price, N. D., Joyce, A. R., et al. (2006) Long-range periodic patterns in microbial genomes indicate significant multi-scale chromosomal organization, PLoS Comput. Biol., 2, e2.

32. Bryant, J. A., Sellars, L. E., Busby, S. J. W., et al. (2014) Chromosome position effects on gene expression in Escherichia coli K-12, Nucleic Acids Res., 42, 11383–11392.

33. Duigou, S. and Boccard, F. (2017) Long range chromosome organization in Escherichia coli: The position of the replication origin defines the non-structured regions and the Right and Left macrodomains, PLoS Genet., 13, e1006758.

34. Herzberg, M., Kaye, I. K., Peti, W., et al. (2006) YdgG (TqsA) controls biofilm formation in Escherichia coli K-12 through autoinducer 2 transport, J. Bacteriol., 188, 587–598.

35. Fuhrer, T., Zampieri, M., Sévin, D. C., et al. (2017) Genomewide landscape of gene-metabolome associations in Escherichia coli, Mol. Syst. Biol., 13, 907.

36. Price, M. N., Wetmore, K. M., Waters, R. J., et al. (2018) Mutant phenotypes for thousands of bacterial genes of unknown function, Nature.

37. Rhee, S. Y. and Mutwil, M. (2014) Towards revealing the functions of all genes in plants, Trends Plant Sci., 19, 212–221.

38. Carrera, J. and Covert, M. W. (2015) Why Build Whole-Cell Models?, Trends Cell Biol., 25, 719–722.

39. Liu, J. K., O’Brien, E. J., Lerman, J. A., et al. (2014) Reconstruction and modeling protein translocation and compartmentalization in Escherichia coli at the genome-scale, BMC Syst. Biol., 8.

40. Seo, S. W., Kim, D., Latif, H., et al. (2014) Deciphering Fur transcriptional regulatory network highlights its complex role beyond iron metabolism in Escherichia coli, Nat. Commun., 5, 4910.

41. Trapnell, C., Williams, B. a., Pertea, G., et al. (2010) Transcript assembly and quantification by RNA-Seq reveals unannotated transcripts and isoform switching during cell differentiation, Nat. Biotechnol., 28, 511–515.

